# Modeling the Structure and DAP5 Binding Site of a Cap-Independent Translational Enhancer mRNA

**DOI:** 10.1101/2023.06.07.542187

**Authors:** Amanda Whittaker, Dixie J. Goss

## Abstract

Cap-independent translation initiation in eukaryotes involves initiation factor (eIF) binding to a transcript’s 5’ untranslated region (UTR). Internal-ribosome-entry-site (IRES)-like cap-independent translation initiation does not require a free 5’ end for eIF binding, as eIFs recruit the ribosome to or near the start codon. For viral mRNA, recruitment usually utilizes RNA structure, such as a pseudoknot. However, for cellular mRNA cap-independent translation, no consensus RNA structures or sequences have yet been identified for eIF binding. Fibroblast-growth factor 9 (FGF-9) is a member of a subset of mRNA that are cap-independently upregulated in breast and colorectal cancer cells using this IRES-like method. Death-associated factor 5 (DAP5), an eIF4GI homolog, binds directly to the FGF-9 5’ UTR to initiate translation. However, the DAP5 binding site within the FGF-9 5’ UTR is unknown. Moreover, DAP5 binds to other, dissimilar 5’ UTRs, some of which need a free 5’ end to stimulate cap-independent translation. We propose that a particular RNA structure involving tertiary folding, rather than a conserved sequence or secondary structure, acts as a DAP5 binding site. Using SHAPE-seq, we modeled the FGF-9 5’ UTR RNA’s complex secondary and tertiary structure in vitro. Further, DAP5 footprinting and toeprinting experiments show DAP5’s preference for one face of this structure. DAP5 binding appears to stabilize a higher-energy RNA fold that frees the 5’ end to solvent and brings the start codon close to the recruited ribosome. Our findings offer a fresh perspective in the hunt for cap-independent translational enhancers. Structural, rather than sequence-specific, eIF binding sites may act as attractive chemotherapeutic targets or as dosage tools for mRNA-based therapies.

## Introduction

Under normoxic conditions, translation initiation occurs in a cap-independent fashion. Eukaryotic initiation factor 4E (eIF4E) recognition of the RNA transcript’s m^7^G cap is necessary for further eukaryotic initiation factor binding and 40S ribosomal recruitment to the transcript (Shatsky *et al*., 2018). However, when oxygen is scarce, 4E-binding protein (4E-BP) sequesters eIF4E and prevents recognition of the m^7^G cap (Braunstein *et al*., 2007). Any ribosomal recruitment to an RNA transcript under hypoxic conditions must therefore proceed in a 4E-independent, or cap-independent, manner: eukaryotic initiation factors bind directly to the 5’ untranslated region (5’ UTR) of the RNA. Eukaryotic viruses and proliferating cancer cells use cap-independent translation to survive and evade host defense mechanisms (Lacerda *et al*., 2017).

Cap-independent translation is believed to initiate via one of two pathways: a cap-independent translational enhancer (CITE)-like mechanism or an internal ribosome entry site (IRES)-like mechanism. Our lab has previously shown that a CITE-like mechanism requires a free 5’ end on the transcript for successful eukaryotic initiation factor binding, ribosomal recruitment and scanning, and cap-independent translation (Haizel *et al*., 2020). Under a CITE-like mechanism of translation initiation, scanning for the start codon by the recruited ribosome is likely. However, a free 5’ end was not shown to be necessary for an IRES-like mechanism, in which ribosomal recruitment to the transcript is thought to occur closer to the start codon. An IRES-like mechanism, therefore, likely requires little to no ribosomal scanning (Haizel *et al*., 2020).

We have previously shown that cap-independent translational upregulation of fibroblast growth factor 9 (FGF-9) occurs in an IRES-like fashion (Haizel *et al*., 2020). FGF-9, an angiogenic growth factor, is translationally upregulated in breast and colorectal cancer cells, where it triggers mitogenesis (Chen *et al*., 2014). Its 5’ UTR contains a regulatory uORF and a predicted IRES (Chen *et al*., 2014). Under normoxic conditions, ribosomes preferentially gather at the uORF start codon; translation of this uORF depresses FGF-9 expression (Chen *et al*., 2014). Under hypoxic conditions, ribosomal presence at the uORF decreases, and ribosomes instead gather at the FGF-9 start codon (Chen *et al*., 2014). Access to the 5’ end of the FGF-9 5’ UTR is not needed for successful cap-independent translation of a downstream luciferase reporter, suggesting the ribosome is recruited away from the m^7^G cap during cellular stress (Haizel *et al*., 2020).

Lack of dependence on this 5’ end invites classification of the FGF-9’s 5’ UTR as a traditional IRES. However, known viral IRES assume complex and functional tertiary structures (Balvay *et al*., 2009), and the secondary and tertiary structures of the FGF-9 5’ UTR are unknown. It has been previously speculated that cellular IRES may assume diverse secondary structures distinct from, yet less complex than, those of viral IRES (Leppek, Das, and Barna, 2018; Komar and Hatzoglou, 2011). The cellular IRES may instead require internal trans-acting factors (ITAFs) to fold the 5’ UTR into a functional structure (Leppek, Das, and Barna, 2018). Indeed, such a mechanism has already been suggested for the FGF-9 5’ UTR, though the exact ITAF that mediates such a change is as yet undiscovered (Chen *et al*., 2014), and the tertiary structure or binding site for such an ITAF has not been identified. Given that cap-independent translation of FGF-9 appears to rely on an IRES-like mechanism, it is therefore crucial to understand what structures, if any, result in successful ribosomal recruitment.

We have previously shown that death associated protein 5 (DAP5), binds the FGF-9 5’ UTR directly; this binding initiates translation of a downstream reporter (Haizel *et al*., 2020). An eIF4GI homolog, DAP5 is overexpressed in proliferating cells (Weingarten-Gabbay *et al*., 2014; Virgili *et al*., 2013). The C-terminal two-thirds of DAP5 are similar to those of eIF4GI; however, the N-terminal region of DAP5 lacks eIF4E and PABP binding sites (Alard *et al*., 2019; Virgili *et al*., 2013). Additionally, DAP5 possesses a longer half-life in the cell under hypoxic conditions as compared to eIF4GI and eIF4GII (Alard *et al*., 2019). DAP5, like eIF4GI, associates with key initiation factors to initiate cap-independent translation; its comparatively long half-life suggests it may substitute for eIF4GI when canonical translation initiation is hindered. Though DAP5 does not unilaterally drive translation initiation when eIF4GI is readily available, it appears to play unique roles in cell survival and immune response, even under normoxic conditions. For example, DAP5 drives Treg cell differentiation in an mTOR/4E-independent, but not cap-independent, manner, via association with *FOXP3* 5’ UTR mRNA (Volta *et al*., 2021).

DAP5 has been previously demonstrated to distinguish between IRES (Weingarten-Gabbay *et al*., 2014). A p53 isoform, Δ*40p53*, has a single IRES within its 5’ UTR and a second IRES within its coding sequence. DAP5 binds both IRES elements with similar affinity, but DAP5 preferentially binds the second IRES *in vivo* (Weingarten-Gabbay *et al*., 2014). How DAP5 distinguishes between these different translational enhancers is unknown, and a universal DAP5 binding site has not yet been identified. Morever, we have previously shown that DAP5 binding does not always stimulate cap-independent translation in an IRES-like manner (Haizel *et al*., 2020). A more complete understanding of DAP5’s role would require characterization of its binding site on one or more 5’ UTRs. Given DAP5’s ability to mediate both cap-dependent and cap-independent translation efficiency, elucidating one or more potential DAP5 binding sites on these transcripts is of broad biological interest.

Computational prediction of cellular, IRES-like elements have not yet produced a putative DAP5 binding site. Neither our group nor others have found significant sequence or secondary structural similarities among 5’ UTRs bound by DAP5 (de la Parra *et al*., 2018), further frustrating efforts to categorize and computationally predict cap-independent translational enhancers. It has previously been suggested that the presence of an upstream uORF, such as FGF-9’s, distinguishes the “true” cellular IRES from other cap-independent translational enhancers (Leppek, Das, and Barna, 2018). However, not all DAP5-dependent transcripts contain uORFs (Shestakova *et al*., 2023). The DAP5 binding site within an IRES-like 5’ UTR is more complex than a single nucleotide sequence or motif which can be identified computationally.

Given the importance of structure to the function of IRES elements, we therefore asked if a complex RNA structural element, rather than a conserved nucleotide sequence or secondary structure, binds DAP5. Here, we present a model of DAP5 binding to the tertiary structure of the FGF-9 5’ UTR mRNA. Using selective 2’-hydroxyl acylation analyzed by primer extension (SHAPE), we found that the 186-nt FGF-9 5’ UTR mRNA likely takes on a stable and highly-structured conformation *in vitro* in the presence of 1 mM magnesium. DAP5 footprinting experiments reveal that DAP5 prefers a single face of this highly-structured 5’ UTR, with the majority of protein-nucleotide contact occurring within the first 93 nucleotides. Additionally, our data suggest DAP5 binding may stabilize a lower-energy, and potentially functional, conformation of the 5’ UTR, where the nt 184-186 AUG codon is more solvent-exposed. This newfound accessibility allows for ribosomal recruitment at or near the start codon. Our proposed model highlights the importance of 5’ UTR RNA structure in DAP5’s function.

## Results

### The FGF-9 5’ UTR

We first sought to determine the FGF-9 5’ UTR secondary and tertiary structure. The FGF-9 5’ UTR was first PCR-amplified from a firefly luciferase reporter construct. In order to ensure the full 5’ UTR was well-represented, a short segment of the firefly luciferase reporter sequence downstream of the 5’ UTR, as well as a short T7 promoter region upstream of the 5’ UTR, were amplified. mRNA was then transcribed *in vitro* using this PCR product as a template. Limiting transcription to the 5’ UTR region prevented base-pairing with the luciferase reporter sequence.

We have previously used SHAPE chemistry as an efficient, time-saving, and accurate method of RNA secondary structure determination (Powell *et al*., 2022). At 37°C, 1-methyl-anhydride-7 (1M7) selectively reacts with nucleotides that are free to move and not associated in base-pairing (Figure 1). If the 2’ oxygen of the ribose sugar is accessible to solvent, 1M7 will react irreversibly to form a 2’ oxygen adduct. Reverse transcriptase is unable to read through this adduct, and the resulting cDNA will stop at the site of 1M7 modification. The resulting cDNA library will therefore show a higher number of reverse transcriptase stops for more flexible nucleotides versus fewer stops for base-paired or inaccessible nucleotides. To distinguish 1M7-mediated stop sites from natural reverse-transcriptase stops, a negative control reaction is run with an equivalent volume of DMSO in place of 1M7 (Mortimer and Weeks, 2007).

**Figure 1.**
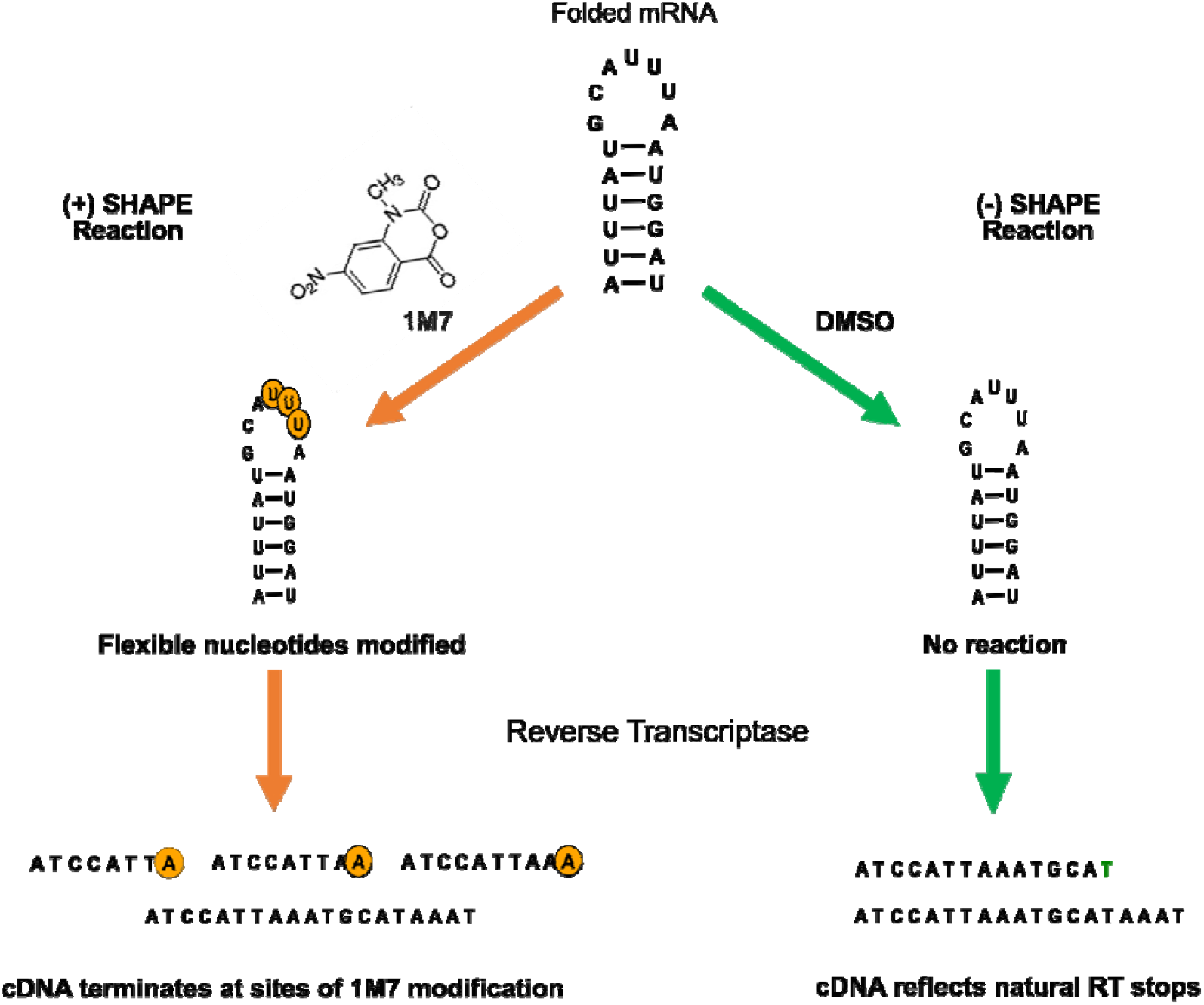
SHAPE Workflow. The transcript of interest is folded in the presence of 1 mM Mg^2+^. Folded RNA is then introduced to 5 mM 1M7 (positive (+) reaction) or an equivalent volume of DMSO as a negative control (negative (−) reaction). 1M7 will form a 2’ oxygen adduct on the ribose sugar of flexible nucleotides. Reverse transcriptase is unable to read through the 2’ oxygen adduct produced by 1M7. The resulting cDNA library terminates at sites of 1M7 modification. In contrast, cDNA produced from DMSO-treated mRNA will only terminate at natural reverse transcriptase stops.

Capillary electrophoresis was used to sequence each cDNA library, and SHAPEFinder software (Vasa *et al*., 2008) calculated 1M7 reactivity per nucleotide. Several repetitions (“runs”) of a folded transcript were compared to determine average 1M7 reactivity (Supplemental Figure 1). Unfolded RNA was not included in our model. These data were fed into RNAStructure software (Reuter and Mathews, 2010) as a folding constraint at 37°C. Predictive folds and associated base-pairing probabilities were generated using a 186-nt region containing the FGF-9 5’ UTR (177 nts) and ending with an AUG (nts 184-186). Finally, 1M7 reactivity was annotated using RNA2Drawer software (Johnson *et al*., 2019).

Using SHAPE chemistry, we found that the FGF-9 5’ UTR takes on a stable, highly-structured conformation (Figure 2A and Supplemental Figure 2). The minimum-free-energy prediction (−73.1 kJ) at 37°C was chosen for analysis (Mortimer and Weeks, 2007). Seven stem loops (SL1 through SL7) were shown to form, with stem loop 1 forming the base of a “pinwheel”-like structure containing stem loops 2 through 5. Stem loop 7 includes the AUG codon (nts 184 - 186) at its base. As expected, 1M7 reactivity is highest for nucleotides in single-stranded or bulge regions. The uORF start codon (nts 24 - 26) was largely unreactive to 1M7.

**Figure 2.**
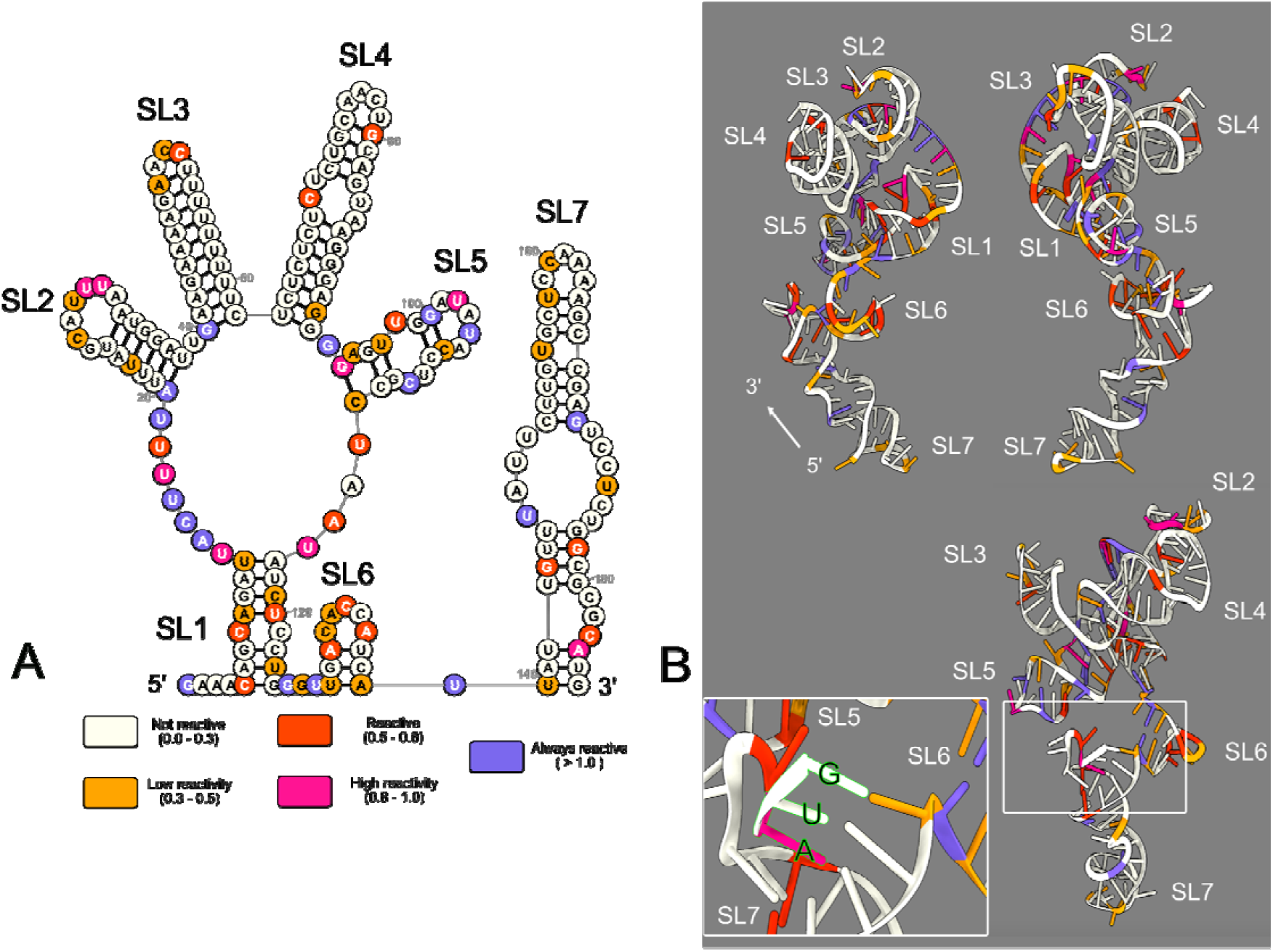
SHAPE-derived 2’ and 3’ structure of the FGF-9 5’ UTR mRNA. Normalized SHAPE data from 3 SHAPE reactions were pooled and fed into RNAStructure software as a binding constraint. The minimum-free-energy predicted structure was selected. Nucleotides were annotated in RNA2Drawer on a scale from 0 (ivory, unreactive to 1M7) to 1.0 and above (medium slate blue, always reactive to 1M7). **A**. The SHAPE-derived 2’ structure of the FGF-9 5’ UTR mRNA. ΔG= -73.1 kJ. Seven stem loops (SL) were predicted to form. The AUG start codon is visible at the base of SL7. **B**. The predicted 3’ structure of the FGF-9 5’ UTR mRNA. The minimum-free-energy tertiary conformation was predicted by RNAComposer using (A) as a folding constraint. USCF ChimeraX was used to annotate 1M7 reactivity. ΔG = -3060 kcal/mol. Stem loops 1-5 aggregate closely around the 5’ end of the mRNA, while stem loops 6 and 7 project away. The AUG start codon, tucked between stem loops 5 and 6, is visualized in an insert (bottom left). SL1 = nucleotides 5-12 and 117-124. SL2 = 20-38. SL3 = 40-62. SL4 = 63-92. SL5 = 94-112. SL6 = 128-138. SL7 = 140 – 186. Start codon: 184 – 186.

RNAStructure’s minimum free-energy fold has been shown to be up to 91% accurate when constrained by 1M7 reactivity (Mortimer and Weeks, 2007). However, RNAStructure cannot predict tertiary conformations. Therefore, we asked how the FGF-9 5’ UTR may sit in three-dimensional space. Our two-dimensional structure was fed into RNAComposer software (Antczak *et al*., 2016; Popenda *et al*., 2012) to generate a minimum-free-energy (−3060 kcal/mol) three-dimensional conformation at 37°C (Figure 2B and Supplemental Movie 1). We annotated 1M7 reactivity on this conformation using UCSF ChimeraX software (Pettersen *et al*., 2020; Goddard *et al*., 2018). Despite its distance from the 5’ end in the two-dimensional structure, stem loop 6 associates closely with stem loops 1 - 5 in a bouquet-like cluster. The 5’ end of the 5’ UTR is tucked within this cluster. A main face, consisting of the stem loop 1 structure, part of the two-dimensional “pinwheel” structure, and stem loop 7, is most accessible to solvent. Interestingly, the FGF-9 start codon is tucked nearest stem loop 6 and is largely inaccessible to 1M7.

### DAP5 Binding

We next asked where the location of DAP5 binding occurs along this highly-structured 5’ UTR. In order to achieve a “ballpark” estimate of where DAP5 binds, we designed a modified version of ribosome toeprinting experiments, nicknamed “DAP5 toeprinting” (Figure 3). Our previous fluorescence binding experiments revealed that DAP5 bound using a single binding site (Haizel *et al*., 2020). The full-length DAP5 peptide was expressed from a plasmid vector and purified as previously described (Haizel *et al*., 2020). To ensure DAP5 was bound to all available transcripts, DAP5 was added to unmodified FGF-9 5’ UTR mRNA at a concentration far above our calculated K_d_ for DAP5 (Haizel *et al*., 2020). Reverse transcriptase was then added directly to the bound mRNA. As reverse transcriptase proceeds along the transcript, it eventually reaches the 3’-most edge of the bound DAP5. The resulting cDNA library overwhelmingly terminates at the edge of DAP5, indicating the edge of the DAP5 binding site. As a negative control, a cDNA library created from unbound, unmodified FGF-9 5’ UTR mRNA was sequenced simultaneously.

**Figure 3.**
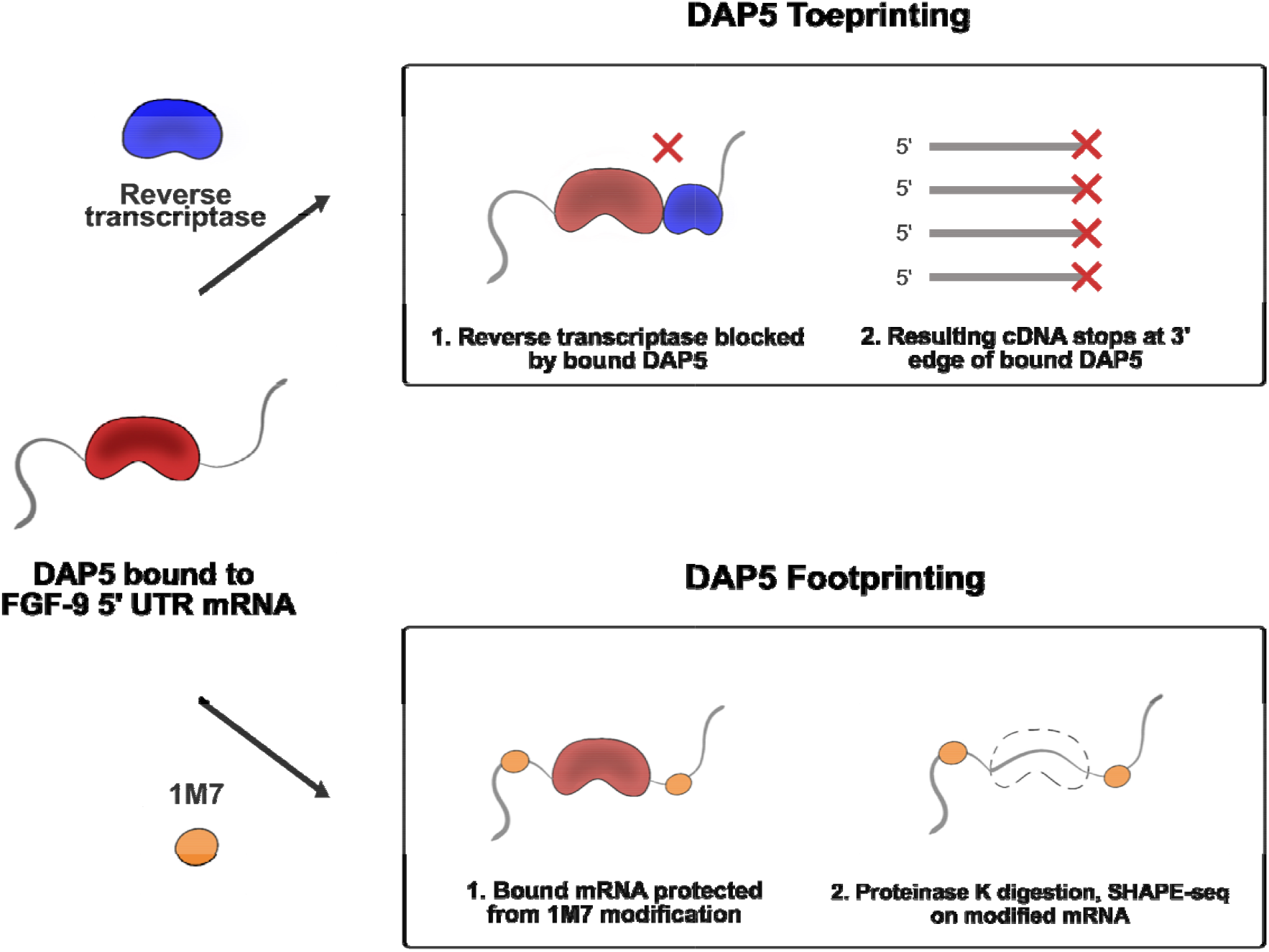
DAP5 Toeprinting versus Footprinting. DAP5 is first bound to folded FGF-9 5’ UTR mRNA. To toeprint (top), reverse transcriptase is introduced directly to bound mRNA. The resulting cDNA library terminates at the 3’-most edge of DAP5. To footprint (bottom), 1M7 is introduced to bound mRNA. DAP5 protects bound nucleotides from 1M7 modification, leading to a localized reduction in SHAPE reactivity. After the 1M7 reaction self-quenches, Proteinase K is introduced to digest DAP5 off of the transcript. cDNA is produced from the modified mRNA and sequenced as described in earlier SHAPE-seq experiments.

We found clear evidence of reverse transcriptase stops almost exclusively at nucleotide 93 of the FGF-9 5’ UTR (Figure 4). This indicates that the bound DAP5 sits among the first four stem loops of the transcript. A large number of cDNA within both the bound and control samples terminated within the short segment of luciferase reporter after the start codon (nucleotide 186), indicating that the full 5’ UTR was well-represented.

**Figure 4.**
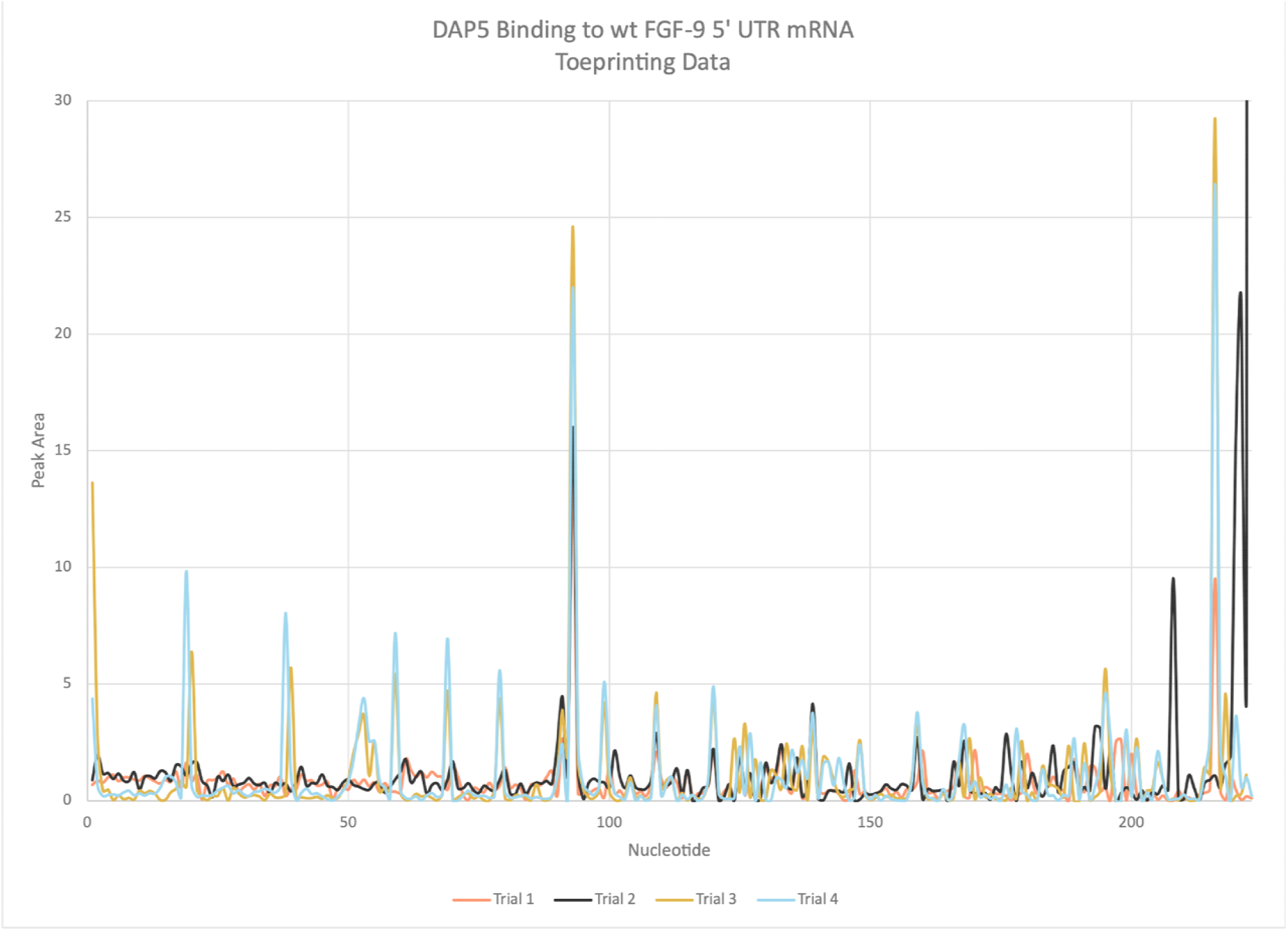
FGF-9 5’ UTR - DAP5 Toeprinting Data. FGF-9 5’ UTR cDNA was reverse-transcribed from DAP5-bound mRNA. n = 4. Higher peak area corresponds to a higher proportion of reverse transcriptase stops in the presence of DAP5. Strong reverse transcriptase stop visible at nucleotide 93. Full-length mRNA peaks are visible between 220 – 240 nucleotides. Start codon: nucleotides 184 – 186.

To better discern the exact nucleotides contacted by DAP5, SHAPE experiments were performed in the presence of DAP5. Unlike toeprinting, DAP5 footprinting involves SHAPE chemistry. DAP5-bound FGF-9 5’ UTR mRNA is modified with 1M7 at 37°C. To ensure all modified RNA is recovered, and to prevent reverse transcriptase stops due to bound protein, DAP5 is digested off of the 1M7-modified RNA using proteinase K (Duncan and Weeks, 2010). Nucleotides that experience a significant decrease in 1M7 reactivity (> 0.3 units) in the presence of bound DAP5 are likely shielded from adduct formation by the protein. “Strong” shielding, where previously accessible nucleotides become inaccessible to 1M7, indicates a strong protein-RNA interaction (Powell *et al*., 2022).

As expected, strong DAP5 contacts were found in the first four stem loops of the FGF-9 5’ UTR (Figure 5A, in green). Nucleotide 93 experienced a sharp decrease in 1M7 reactivity in the presence of DAP5. Smaller decreases in 1M7 reactivity are visible at individual nucleotides throughout the FGF-9 5’ UTR transcript (Figure 5A). These decreases appear scattered across our proposed secondary structure (Figure 5B). However, these DAP5 contact points coalesce along a single face of our proposed tertiary structure (Figure 5C and Supplemental Movie 2). Much of this contact is centered around the start codon (magenta) at the end of SL7.

**Figure 5.**
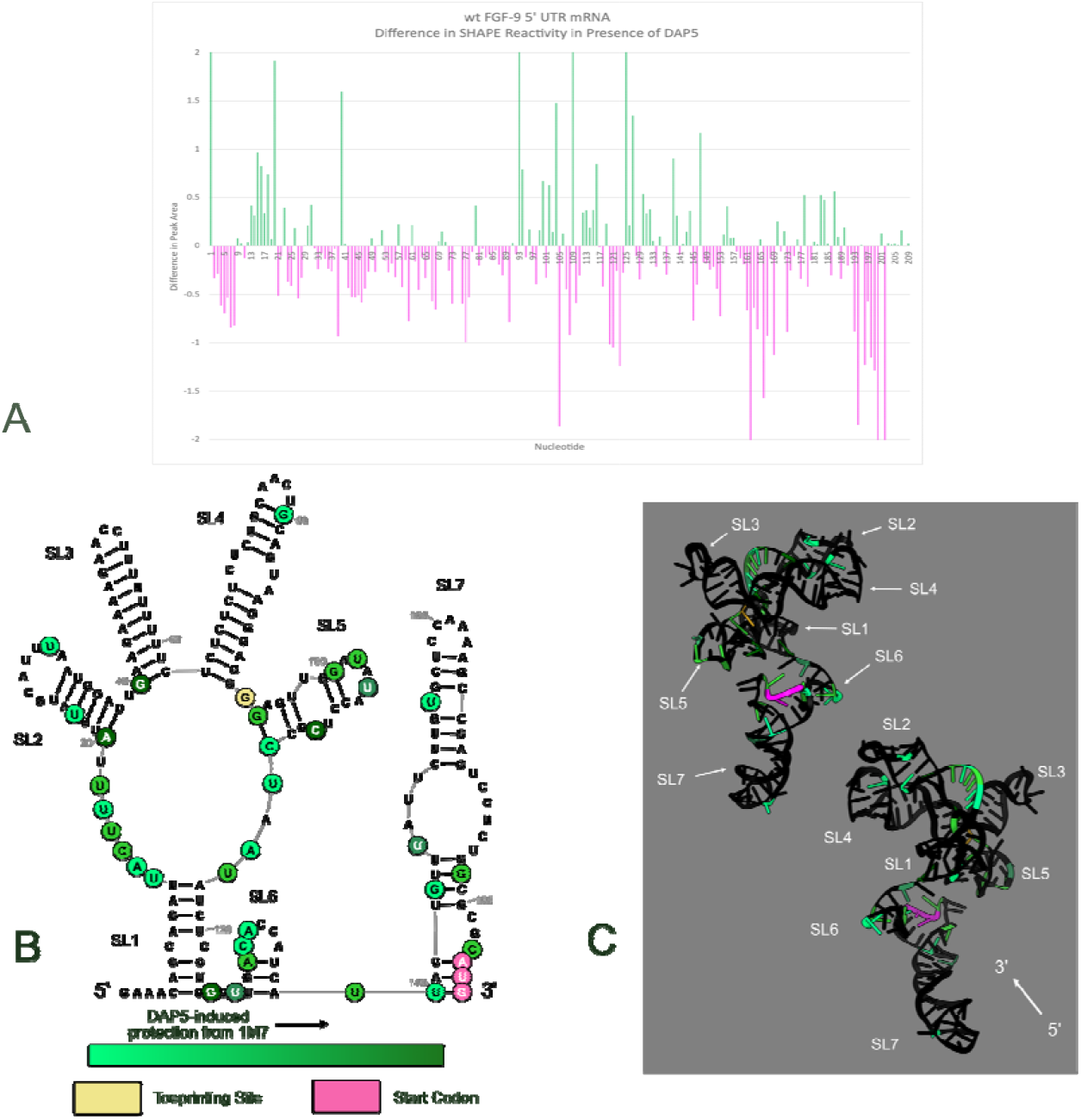
FGF-9 5’ UTR - DAP5 Footprinting Data. 1M7 reactions were performed on DAP5-bound FGF-9 5’ UTR mRNA. n = 2. **A**. Changes in 1M7 reactivity in the presence of DAP5. Normalized 1M7 reactivity in the presence of DAP5 was subtracted from normalized 1M7 reactivity in the absence of DAP5. A decrease in reactivity (green) indicates possible contact with DAP5. An increase in reactivity (magenta) indicates increased flexibility in the presence of DAP5. Start codon: nt 184 – 186. Full length of mRNA = 209 nucleotides. **B**. DAP5-induced protection from 1M7 annotated across the predicted secondary structure of the FGF-9 5’ UTR mRNA. Decreased 1M7 reactivity is annotated in a green gradient, with darker green = sharper decrease in reactivity. The toeprinting site (nt 93) from Fig. 4 is annotated in tan. The start codon is annotated in pink. **C**. DAP5-induced protection from 1M7 annotated on the predicted tertiary structure of the folded FGF-9 5’ UTR mRNA. All other nucleotides are annotated in black.

Interestingly, we noticed a significant increase in 1M7 reactivity (pink) for some nucleotides (Figure 5A, in pink). Increased accessibility to 1M7 indicates a change in secondary or tertiary structure: for example, a previously base-paired nucleotide may become single-stranded. In particular, the short luciferase reporter region after the start codon (nt 187+) became more accessible to 1M7 modification in the presence of DAP5. SL3 and SL4 (nts 40 - 90) were slightly more reactive to 1M7, indicating increased flexibility.

## Discussion

We sought to characterize the FGF-9 5’ UTR and identify its DAP5 binding site. Our SHAPE-derived model of the FGF-9 5’ UTR mRNA includes a complex secondary and tertiary structure. Further, DAP5 binds to a single face of the FGF-9 5’ UTR, reaching as far as nucleotide 93. While DAP5 contact appears scattered across our secondary structure model, these localized contacts coalesce into a “binding surface” across our tertiary model. We therefore propose that the DAP5 binding site is not a localized sequence or single nucleotide, but a preference for one face of the 5’ UTR.

FGF-9’s tertiary structure does not contain the structural trademarks of the viral IRES. Though our toeprinting site occurs in a guanine-rich region, no other guanine-rich sequence is available for G-quadruplex formation. We likewise did not find evidence of pseudoknot formation. The pinwheel structure formed by stem loops 1 - 5 is reminiscent of stem loop K of the EMCV IRES, where eIF4GI binds (Balvay *et al*., 2009). Though FGF-9 has long been speculated to be an IRES, its complex structure mimics, but does not conform to, the structure of known viral IRES. Nevertheless, the structural complexity of the FGF-9 5’ UTR indicates that the structure of a cellular IRES may be, as previously speculated, distinct from that of a viral IRES (Komar and Hatzoglou, 2011). We will continue to refer to the FGF-9 5’ UTR as “IRES-like.”

Unexpectedly, DAP5 binding seems to rearrange the FGF-9 5’ UTR and stabilize a lower-energy fold (Figure 6 and Supplemental Movie 3). DAP5 associates with eIF4A, an ATP-dependent RNA helicase, but our findings suggest DAP5 itself may influence local RNA structure in the absence of eIF4A (Virgili *et al*., 2013). 1M7 reactivity in the presence of DAP5 was used as an RNAStructure folding constraint. Our seven-stem-loop model resolved into four modified stem loops (MSL) with a slightly higher free energy (−55.0 kJ) than our original fold (Figure 6A and Supplemental Figure 3). The uORF start codon (nts 24 - 27, within MSL1) became highly reactive to 1M7. When this RNAStructure fold was pushed into RNAComposer, these modified stem loops coalesced into a flat, triangular structure. The 5’ end, now single-stranded, is exposed to solvent (Figure 6B). DAP5 contacts cluster around the toeprinting site nearest the start codon (Figure 6C and Supplemental Movie 4). The 5’ end, however, lacks DAP5 contacts. This finding is consistent with our earlier studies (Haizel *et al*., 2020) showing that a 5’-most stem loop did not inhibit the cap-independent translation of a downstream reporter sequence.

**Figure 6.**
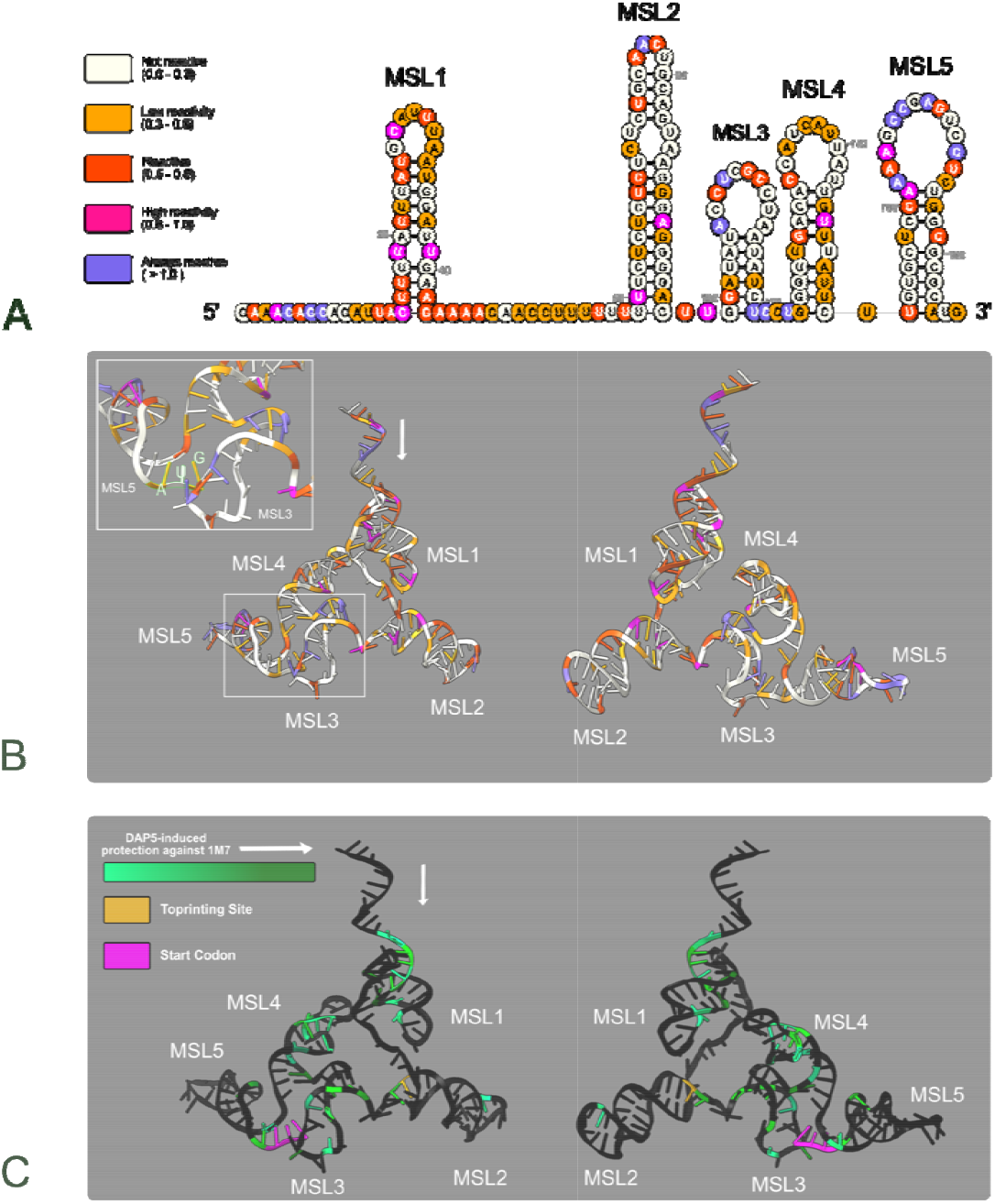
Predicted structural effect of DAP5 Binding on the FGF-9 5’ UTR. The 1M7 reactivity of the FGF-9 5’ UTR mRNA in the presence of DAP5 was fed into structural prediction software. n=2. **A**. Lowest-free-energy conformation of the bound FGF-9 5’ UTR as predicted in RNAStructure. Five modified stem loops (MSL) were predicted. ΔG = -55.7 kJ. Nucleotides are colored for 1M7 reactivity as previously described (Fig. 2). **B**. The lowest-free energy tertiary conformation of (A) predicted by RNAComposer. ΔG= -3530 kcal. The start codon visualized in top left insert. **C**. DAP5 footprinting data (Fig. 5) annotated on (B). 1M7 reactivity decreases in the presence of DAP5 are annotated in a green gradient, with darker green = sharper decrease. Magenta = start codon. Toeprinting site (nt 93) = goldenrod. All other nucleotides annotated in black. MSL1 = nucleotides 15-43. MSL2 = 59-96. MSL3 = 99-120. MSL4 = 124-151. MSL5 = 153 – 184.

Given that DAP5 does not need a free 5’ end to drive cap-independent translation initiation of FGF-9, it follows that DAP5 would interact minimally, if at all, with this 5’ end. Additional contacts near the start codon suggest that DAP5 may recruit the ribosome directly to the start codon, and that this codon is more exposed when DAP5 is bound.

It has been previously suggested that, as with other cellular IRES, an unknown ITAF changes FGF-9’s secondary structure into a cap-independent translational enhancer under hypoxia (Chen *et al*., 2014). Our data suggest DAP5 itself is one such ITAF. Indeed, DAP5 has already been named as an ITAF for other IRES-like translational enhancers, such as Δ40p53 (Marques, Lacerda, and Romão, 2022). We have shown here that DAP5 binding frees the FGF-9 AUG from SL7, putting it within reach of DAP5 and, potentially, the recruited ribosome (Fig. 6C). This agrees with previous findings showing ribosomal preference for the FGF-9 start codon under hypoxia (Chen *et al*., 2014).

Overall, we find that a DAP5-mediated change agrees with our group’s previous IRES-like model (Haizel *et al*., 2020). Here, we provide strong evidence that DAP5 acts as an ITAF to promote rearrangement of the 5’ UTR and promote translation of the FGF-9 mRNA at the FGF-9 AUG, rather than the uORF. Integrating our RNA structural data into computational predictions allowed us to characterize this IRES-like element. This model provides insight into recognition features of the RNA, which offer potential for use in designing novel chemotherapeutic targets and may aid in identifying other complex, IRES-like RNA structural elements.

## Acknowledgements

This project was funded by NIH RO1 GM 128239 and NSF MCB 1902054.

The authors would wish to thank Dr. Usha Bhardwaj, Dr. Solomon Haizel, and Dr. Paul Powell for gifting their plasmid constructs, primers, and knowledge of SHAPE.

Open-source software was crucial for this publication. The authors thank the Laederach lab (UNC) for their MATLAB CEQ alignment suite. Additional thanks are in order for the Weeks lab and Giddings lab (both UNC Chapel Hill) for providing us with SHAPEFinder software. Thank you to the Mathews Lab (URMC) for RNAStructure, used for our 2D structural predictions. 2D structures were annotated thanks to RNA2Composer, developed by the Simon Lab (UMD) in collaboration with the NCI. We are grateful to the RNAComposer team (Polish Academy of Sciences, Poznan University of Technology) for their 3D modeling software. 3D structures were annotated thanks to UCSF ChimeraX, developed by the Resource for Biocomputing, Visualization, and Informatics at the University of California, San Francisco.

## Methods

### Isolation of FGF-9 5’ UTR Region

Plasmid reporter constructs containing a T7 promoter region and the 5’ UTR sequence of FGF-9 (177 nts, GenBank Accession Number: AY682094.1) directly upstream of a firefly luciferase coding region were a generous gift from Dr. Solomon Haizel (Haizel *et al*., 2020). To simplify later SHAPE-seq analysis, the T7 promoter, FGF-9 5’ UTR sequence, and the first 75 nucleotides of the firefly luciferase reporter were PCR-amplified using the ThermoFisher PCR Phusion High-Fidelity Kit. The resulting PCR product was purified into nuclease-free water using the Zymo Oligo Clean and Concentrator kit. The concentration was verified using a NanoDrop UV/Vis spectrophotometer, and integrity was confirmed using a native 1.5% w/v agarose gel.

### FGF-9 RNA Transcription

RNA was *in vitro* transcribed overnight from the isolated FGF-9 5’ UTR region using the NEB HiScribe T7 High Yield RNA Synthesis Kit. To prevent RNA degradation, 1 μL of a 1.6 units/μL ThermoScientific RiboLock RNase inhibitor solution was included. The 75 nucleotides of the firefly luciferase reporter and a short 20-nucleotide region upstream of the FGF-9 5’ UTR were retained in the resulting transcript to ensure the entire FGF-9 5’ UTR was well-represented in later SHAPE data. After DNase digestion, RNA was purified into nuclease-free water using the Zymo Oligo Clean and Concentrator kit. As with the PCR product, RNA concentration was verified using a NanoDrop UV/Vis spectrophotometer, and RNA integrity was confirmed using a native 1.5% w/v agarose gel.

### SHAPE-seq

50 mM 1-methyl-7-nitroisatoic anhydride (1M7) and pure DMSO stock solutions were pre-heated to 37°C prior to SHAPE reactions. 20 pmol FGF-9 5’ UTR mRNA was suspended in 20 mM HEPES buffer (pH = 7.5) with 100 mM KCl. As described previously (Haizel *et al*., 2020), nascent RNA secondary and tertiary structure was disrupted by heating to 90°C over 2 minutes, followed by slow cooling at room temperature for 1 hour. To refold RNA, MgCl_2_ was added to a final concentration of 1 mM Mg^2+^ before incubation on ice for 1 hour (Haizel *et al*., 2020).

SHAPE-seq reactions were executed according to existing literature (Powell *et al*., 2022; Rice e*t al*., 2014). For positive (+) SHAPE reactions, pre-heated 1M7 stock was added to folded RNA sample to a final concentration of 5 mM. For negative control (−) reactions, an equivalent volume of pre-heated DMSO was added to sample in lieu of 1M7. Both solutions were incubated at 37°C for 70 seconds, equivalent to five 1M7 half-lives (Mortimer and Weeks, 2007). Sample tubes were wrapped in tinfoil to protect 1M7-modified RNA from light. RNA was purified into 9 μL nuclease-free water using the Zymo Oligo Clean and Concentrator kit.

A cyanine-5 (cy5)-labeled cDNA primer corresponding to part of the firefly luciferase reporter region preserved in the amplified DNA sequence was a generous gift from Dr. Paul Powell (Supplemental Table 1). Reverse transcription of (+) and (−) SHAPE samples was executed using the Invitrogen SuperScript III kit as previously described (Powell *et al*., 2022). 10 pmols of cy5-labeled cDNA primer, 1 μL of RiboLock RNase inhibitor, 2 μL of NEB10 μM dNTP mix, 1 μL of 0.1 mM DTT, 1 μL nuclease-free water, and 4 μL Invitrogen 5x first-strand buffer were added to SHAPE samples. CTP-sequencing samples were created using unreacted FGF-9 5’ UTR mRNA and 1 μL ddGTP in lieu of nuclease-free water. For UTP-sequencing samples, ddATP was used in lieu of ddGTP. To allow primer annealing, all samples were incubated at 65°C for 5 minutes before slow cooling to 55°C. 1 μL Invitrogen SuperScript III reverse-transcriptase was added before incubation at 55°C for 40 minutes.

Remaining RNA was digested via addition of 2 μL 2M NaOH and incubation at 95°C for 3 minutes, followed by immediate neutralization with 2 μL 2M HCl. cDNA was recovered via ethanol precipitation at 4°C overnight (Powell *et al*., 2022). cDNA was sequenced using capillary electrophoresis on the Beckman GenomeLab GeXP Genetic Analysis System using custom parameters (Powell *et al*., 2022; Mitra *et al*., 2008).

### 5’ UTR Structural Prediction

Simultaneous capillary electrophoresis runs were first aligned according to their size standards using the MATLAB CEQ alignment suite of software developed by the Laederach Lab at UNC Chapel Hill (available at https://ribosnitch.bio.unc.edu/software/). .shape files were then created using one (+) trace, one (−) trace, one CTP-seq trace, and one UTP-seq trace. SHAPEFinder software, developed by the Giddings Lab at UNC Chapel Hill (Vasa *et al*., 2008), was used to determine 1M7 reactivity per nucleotide. The difference in peak area between (+) capillary electrophoresis trace and the (−) capillary electrophoresis trace defined the degree of 1M7 SHAPE reactivity. A higher difference in peak area corresponds to a more frequent reverse-transcriptase stop in the (+) sample and, therefore, a more reactive nucleotide.

Individual SHAPE runs were normalized against themselves using the “simple” normalization method described in Low and Weeks, 2010. The highest 2% of reactivities across all nucleotides were first temporarily removed from the data pool. The following top 8% of reactivities were averaged against each other to create a normalization factor. The removed reactivity values were then re-added to the data pool, and all reactivities were divided by the calculated normalization factor. This resulted in a normalized 1M7 reactivity value for each nucleotide ranging between 0 (not reactive to 1M7) to > 1.5 (extremely reactive to 1M7).

In order to determine the average 1M7 reactivity for each nucleotide, multiple SHAPE runs were normalized against each other using the “box-plot” normalization method (Rice *et al*., 2014; Low and Weeks, 2010). Normalized 1M7 reactivity values were created for each individual nucleotide across multiple runs. The 25^th^ and 75^th^ quartile, as well as the Q value, was calculated for each nucleotide’s data pool, as well as the Q value. If a datum was greater than 1.5Q, it was labeled as an outlier. However, if 1.5Q < 90^th^ percentile, data greater than the 90^th^ percentile were labeled as outliers instead. Chosen outliers were summarily removed from the data pool. Remaining data points were averaged and fed into RNAStructure software as a folding constraint at 37°C (Reuter and Mathews, 2010). The RNAStructure output was annotated for SHAPE reactivity and base-pairing probabilities using RNA2Drawer software (Johnson *et al*., 2019).

RNAComposer software (Antczak *et al*., 2016; Popenda *et al*., 2012) was used to generate 3D structural predictions. The MFE structure, generated by RNAStructure, was fed into RNAComposer. The MFE conformation at 37°C was then generated. Visualization and annotation of this 3D structure, as well as a “spin video” showing the structure throughout a 360° turn, were made using UCSF Chimera X software (Pettersen *et al*., 2020; Goddard *et al*., 2018).

### DAP5 Isolation

A plasmid construct, including the DAP5 coding sequence followed by an N-terminal 6x histidine tag, was a generous gift from Dr. Solomon Haizel. A TEV protease cut site preceding the 6x histidine tag was previously introduced using the Q5 Site-Directed Mutagenesis Kit from NEB and the GeneJET Plasmid Miniprep Kit from ThermoFisher. DAP5 was expressed in *Escherichia coli* BL21-CodonPlus (DE3)-RIL cells from Agilent and purified as previously described (Haizel *et al*., 2020). DAP5 was first purified from bacterial cell lysate using a 5 mL His-Trap HP (Ni-NTA) column from GE Healthcare Life Sciences. To cleave the N-terminal 6x histidine tag, purified DAP5 was dialyzed overnight at 4°C against storage buffer (20 mM HEPES (pH 7.5), 200 mM KCl, 10% glycerol, 20 mM BME) in the presence of TEV protease. Final purification and concentration of the untagged DAP5 was performed using a 1 ml HiTrap heparin HP column from GE Healthcare Life Sciences. Elution fraction quality was verified using a 10% SDS-PAGE gel; chosen elution fractions (> 95% purity) were pooled and distributed into uniform aliquots. Protein concentration was then determined using ThermoScientific’s Coomassie Blue protein assay reagent. Aliquots were stored at -80°C. To ensure protein integrity, used aliquots were discarded after 3 freeze-thaw cycles.

### DAP5 Toeprinting

20 pmol FGF-9 5’ UTR RNA was first folded in the presence of magnesium. To ensure the majority of RNA was bound by protein, purified DAP5 was introduced to a final concentration of 300 μM, well above our group’s determined K_d_ for DAP5 binding to the FGF-9 5’ UTR (Haizel *et al*., 2020; Tijerina *et al*., 2007). To match SHAPE solutions, the final volume of these protein-RNA solutions was 9 μL. As a negative control, 9 μL of a 20 pmol FGF-9 5’ UTR RNA solution was run in parallel. Protein-RNA solutions were incubated at room temperature for 10 minutes. 10 pmols cy5-labeled cDNA primer, 1 μL of RiboLock RNase inhibitor, 2 μL of NEB10 uM dNTP mix, 1 μL of 0.1 mM DTT, 1 μL nuclease-free water, and 4 μL Invitrogen 5x first-strand buffer were added directly to protein-RNA solutions. Parallel CTP-seq and UTP-seq reactions were run with unbound FGF-9 mRNA and ddNTPs. Reverse-transcription, cDNA purification, and sequencing via capillary electrophoresis were performed as described for SHAPE samples (see SHAPE-seq). Peak height in the presence of DAP5 was determined using SHAPEFinder (Vasa *et al*., 2008). The “box-plot” method was then used for data normalization (Rice *et al*., 2014; Low and Weeks, 2010).

### DAP5 Footprinting

1M7 and DMSO were pre-heated to 37°C. As with toeprinting experiments, 300 μM DAP5 was bound to folded 20 pmol FGF-9 5’ UTR RNA at room temperature for 10 minutes. 5 mM 1M7 or an equivalent volume of DMSO were added to protein-RNA solutions, followed by incubation at 37°C for 70 seconds. To digest DAP5 off of RNA and ensure all modified RNA was fully recovered, 60 μg Proteinase K was added, followed by incubation at 37°C for 10 minutes (Duncan and Weeks, 2008). Sample tubes were again wrapped in tinfoil to protect 1M7-modified RNA from light. 1M7-modified RNA was purified into 9 μL nuclease-free water using the Zymo Oligo Clean and Concentrator kit. Reverse-transcription, cDNA isolation, sequencing, SHAPEFinder analysis, and data normalization were performed as described with SHAPE samples. To determine changes in 1M7 reactivity, normalized and averaged 1M7 reactivity in the presence of DAP5 was subtracted from normalized and averaged 1M7 reactivity in the absence of DAP5. Normalized and averaged 1M7 reactivity in the presence of DAP5 was used as a folding constraint at 37°C in RNAStructure, and the MFE fold was chosen. The RNAStructure 2D MFE fold was used in RNAComposer to generate the 3D MFE fold at 37°C.

## Notes

### Competing Interest Statement

The authors have declared no competing interest.

