## Supplemental Figures for "Modeling the Structure and DAP5 Binding Site of a Cap-Independent Translational Enhancer mRNA": DAP5_FGF9_SupplementalFigures_Biorxiv.pptx

### Slide 1
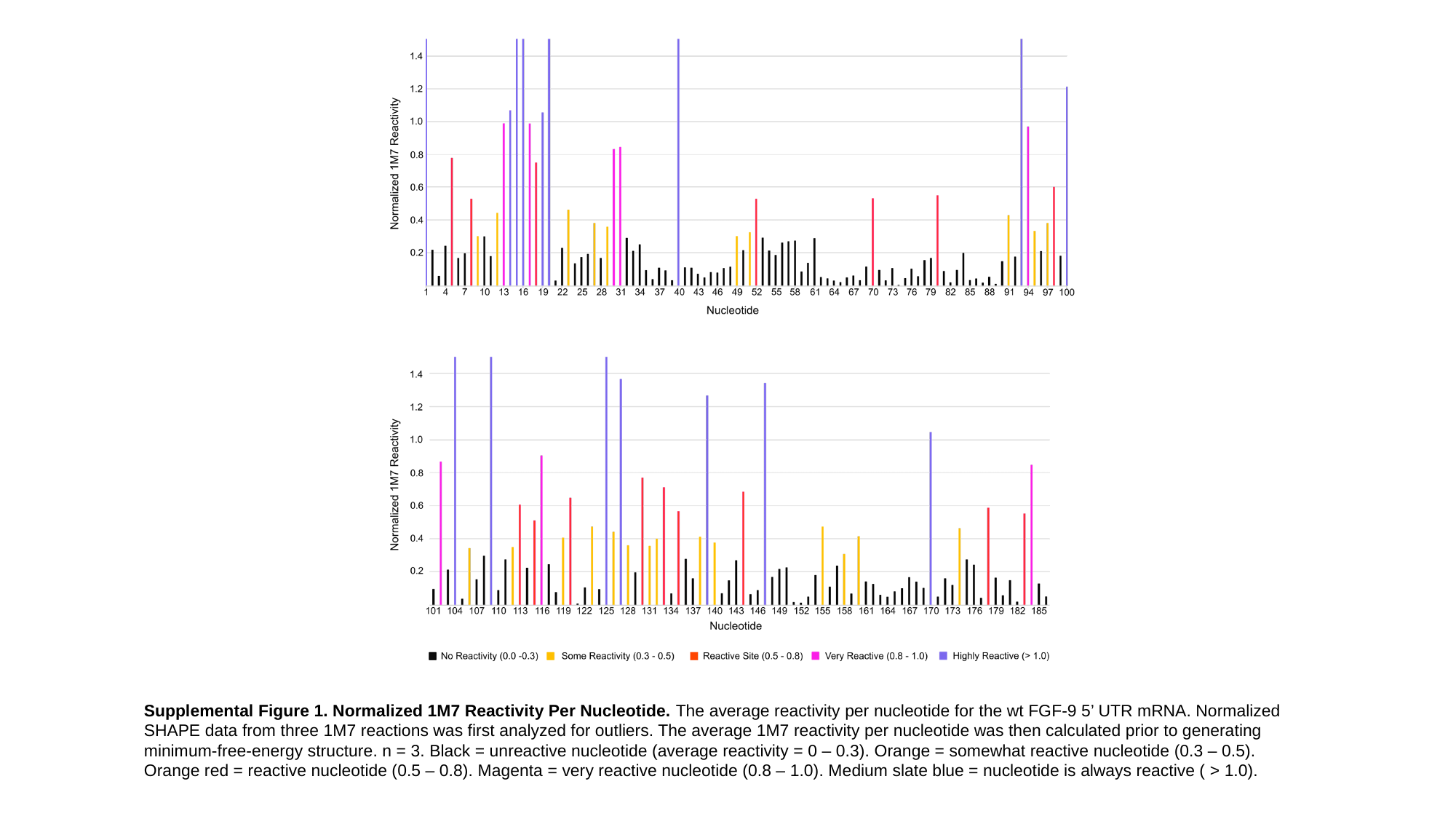

Supplemental Figure 1. Normalized 1M7 Reactivity Per Nucleotide. The average reactivity per nucleotide for the wt FGF-9 5’ UTR mRNA. Normalized SHAPE data from three 1M7 reactions was first analyzed for outliers. The average 1M7 reactivity per nucleotide was then calculated prior to generating minimum-free-energy structure. n = 3. Black = unreactive nucleotide (average reactivity = 0 – 0.3). Orange = somewhat reactive nucleotide (0.3 – 0.5). Orange red = reactive nucleotide (0.5 – 0.8). Magenta = very reactive nucleotide (0.8 – 1.0). Medium slate blue = nucleotide is always reactive ( > 1.0).

### Slide 2
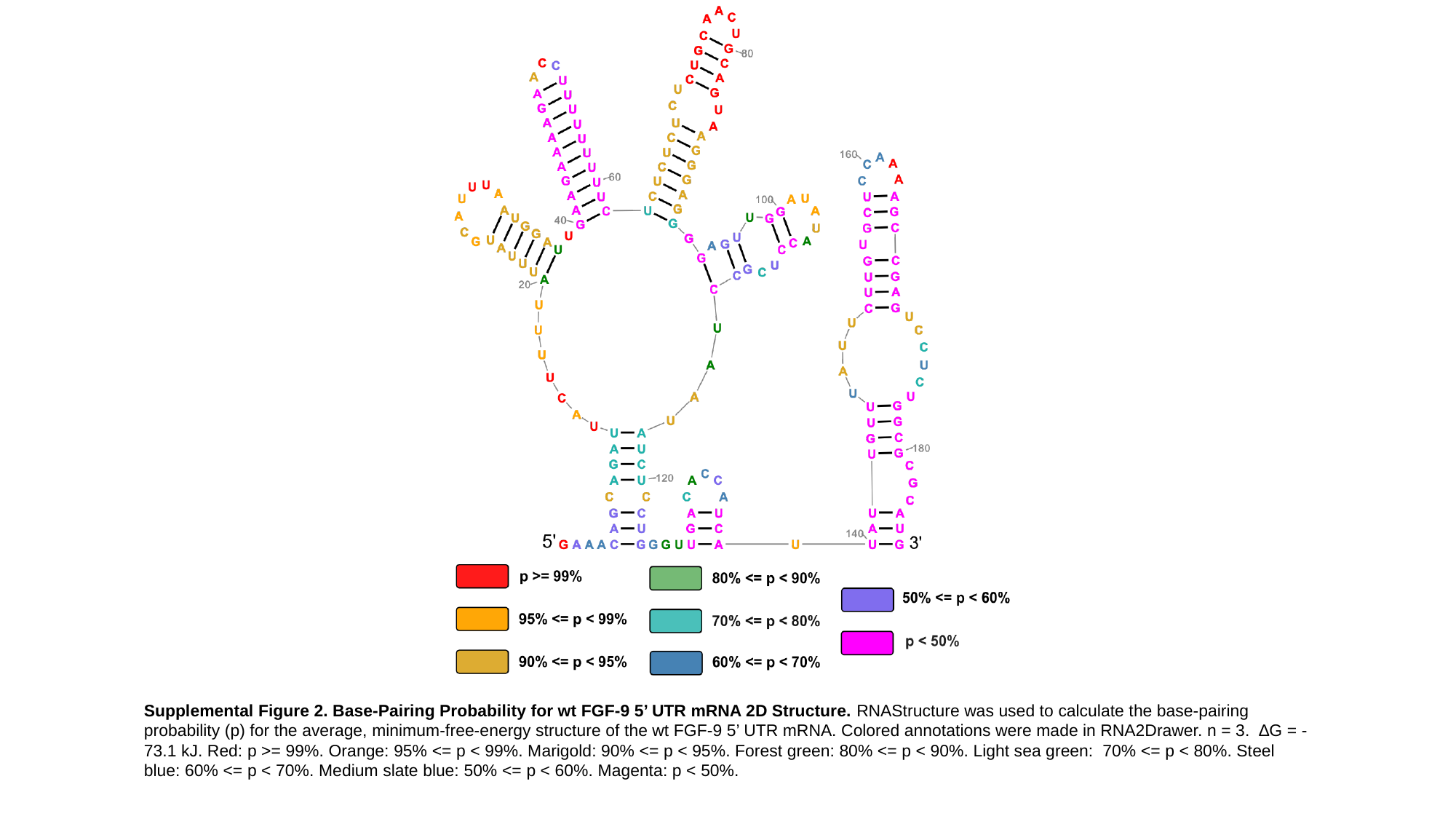

Supplemental Figure 2. Base-Pairing Probability for wt FGF-9 5’ UTR mRNA 2D Structure. RNAStructure was used to calculate the base-pairing probability (p) for the average, minimum-free-energy structure of the wt FGF-9 5’ UTR mRNA. Colored annotations were made in RNA2Drawer. n = 3. ∆G = -73.1 kJ. Red: p >= 99%. Orange: 95% <= p < 99%. Marigold: 90% <= p < 95%. Forest green: 80% <= p < 90%. Light sea green: 70% <= p < 80%. Steel blue: 60% <= p < 70%. Medium slate blue: 50% <= p < 60%. Magenta: p < 50%.

### Slide 3
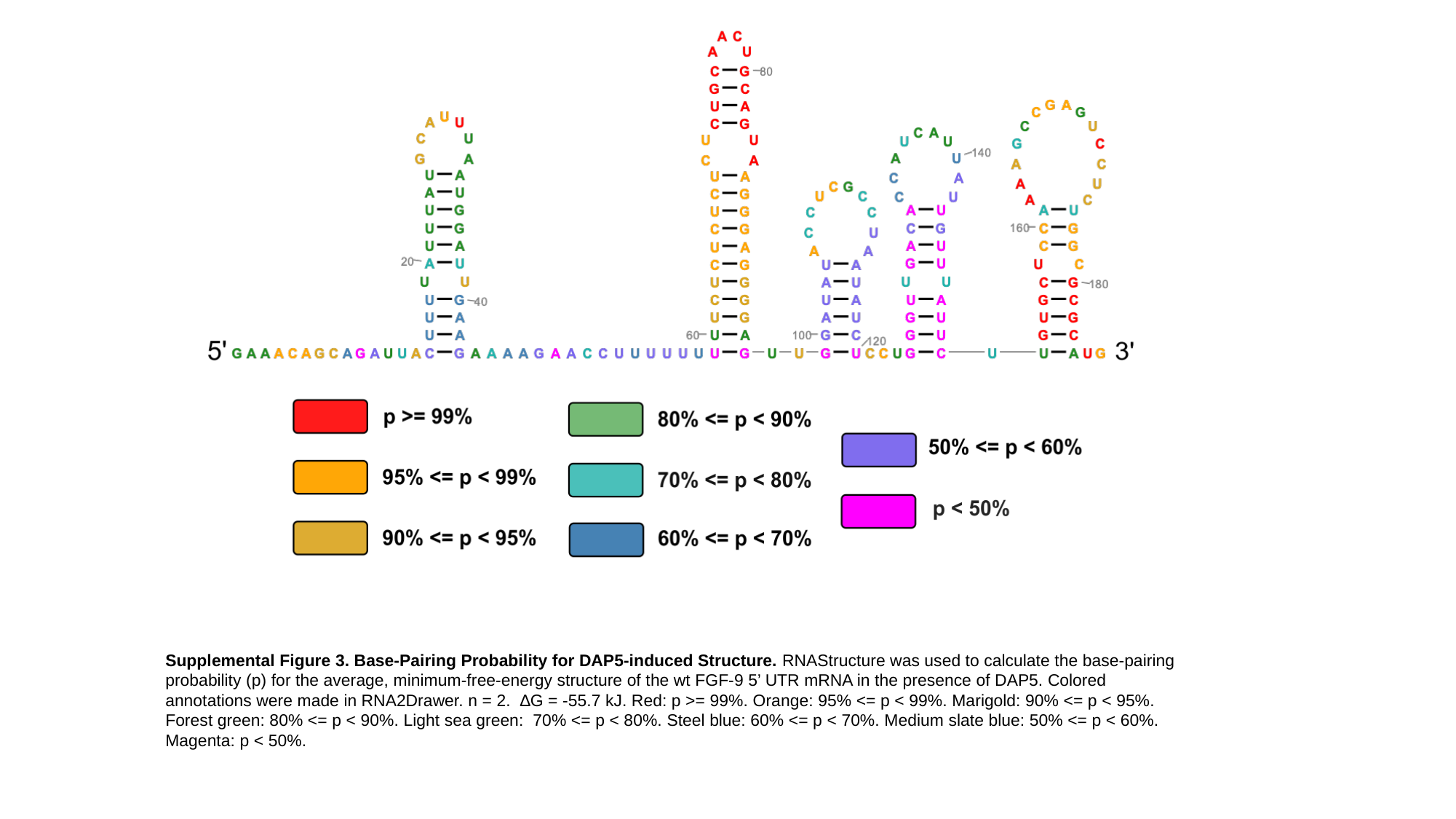

Supplemental Figure 3. Base-Pairing Probability for DAP5-induced Structure. RNAStructure was used to calculate the base-pairing probability (p) for the average, minimum-free-energy structure of the wt FGF-9 5’ UTR mRNA in the presence of DAP5. Colored annotations were made in RNA2Drawer. n = 2. ∆G = -55.7 kJ. Red: p >= 99%. Orange: 95% <= p < 99%. Marigold: 90% <= p < 95%. Forest green: 80% <= p < 90%. Light sea green: 70% <= p < 80%. Steel blue: 60% <= p < 70%. Medium slate blue: 50% <= p < 60%. Magenta: p < 50%.

### Slide 4
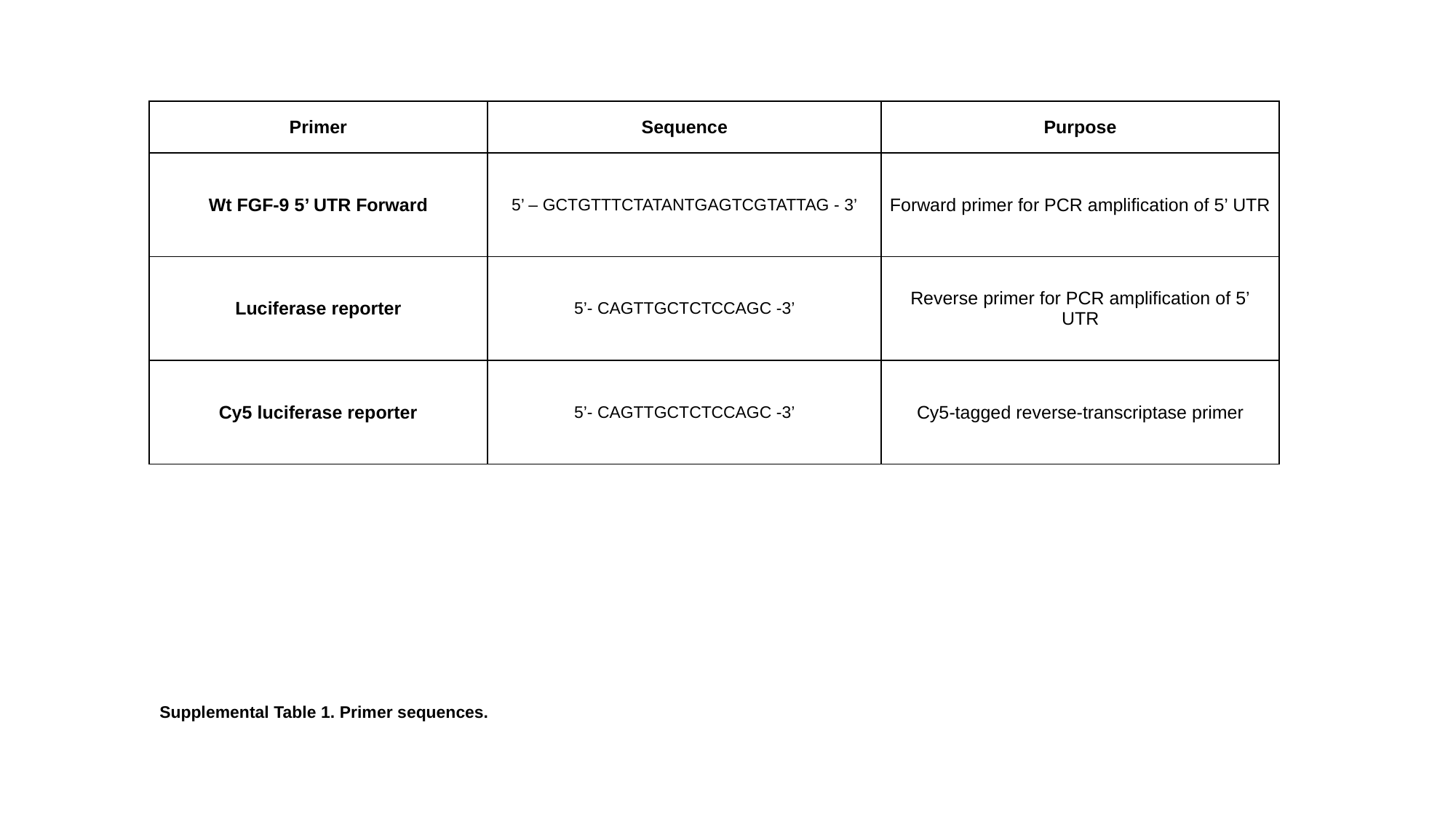

| Primer | Sequence | Purpose |
| --- | --- | --- |
| Wt FGF-9 5’ UTR Forward | 5’ – GCTGTTTCTATANTGAGTCGTATTAG - 3’ | Forward primer for PCR amplification of 5’ UTR |
| Luciferase reporter | 5’- CAGTTGCTCTCCAGC -3’ | Reverse primer for PCR amplification of 5’ UTR |
| Cy5 luciferase reporter | 5’- CAGTTGCTCTCCAGC -3’ | Cy5-tagged reverse-transcriptase primer |
Supplemental Table 1. Primer sequences.

### Slide 5
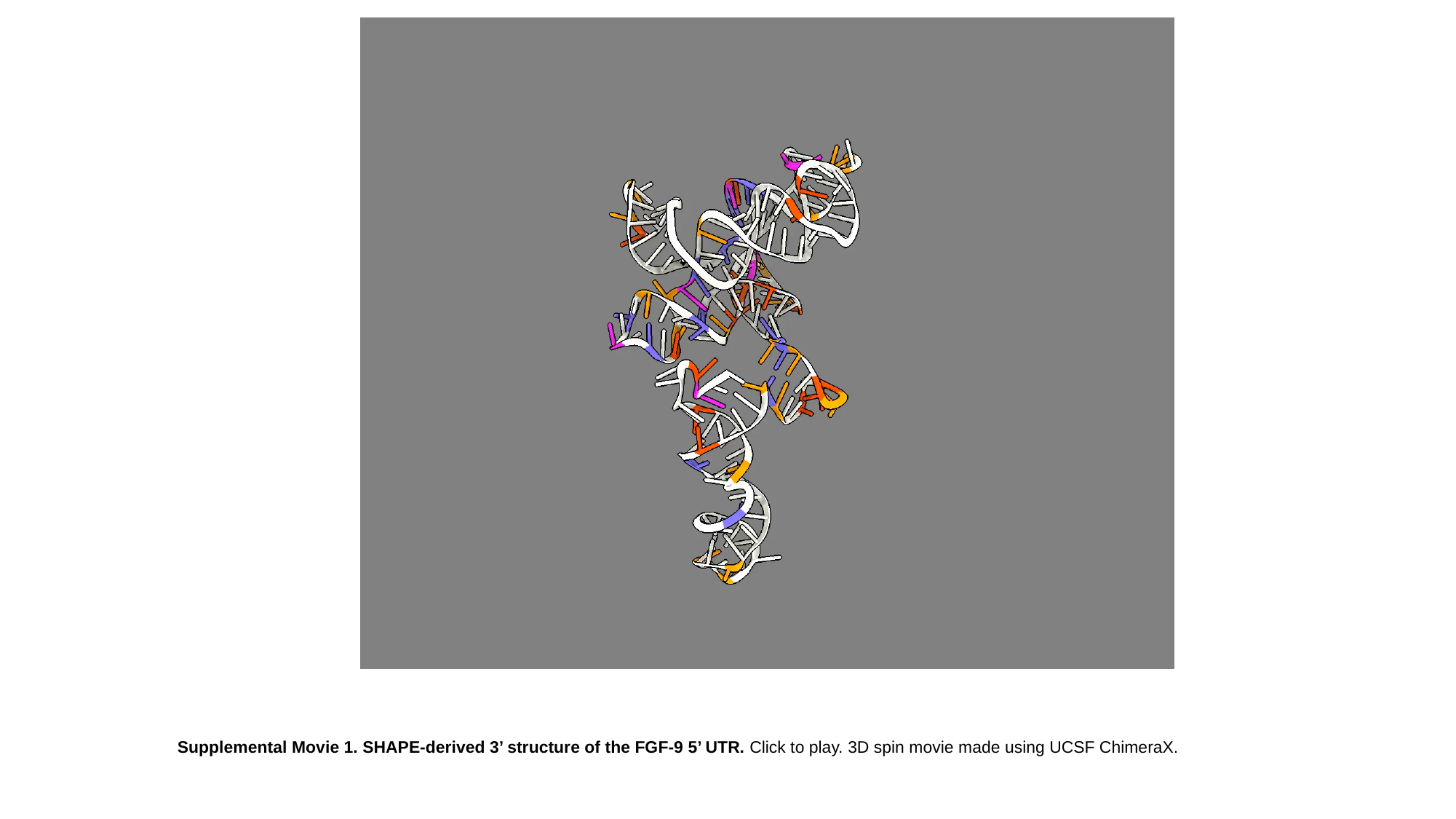

Supplemental Movie 1. SHAPE-derived 3’ structure of the FGF-9 5’ UTR. Click to play. 3D spin movie made using UCSF ChimeraX.

### Slide 6
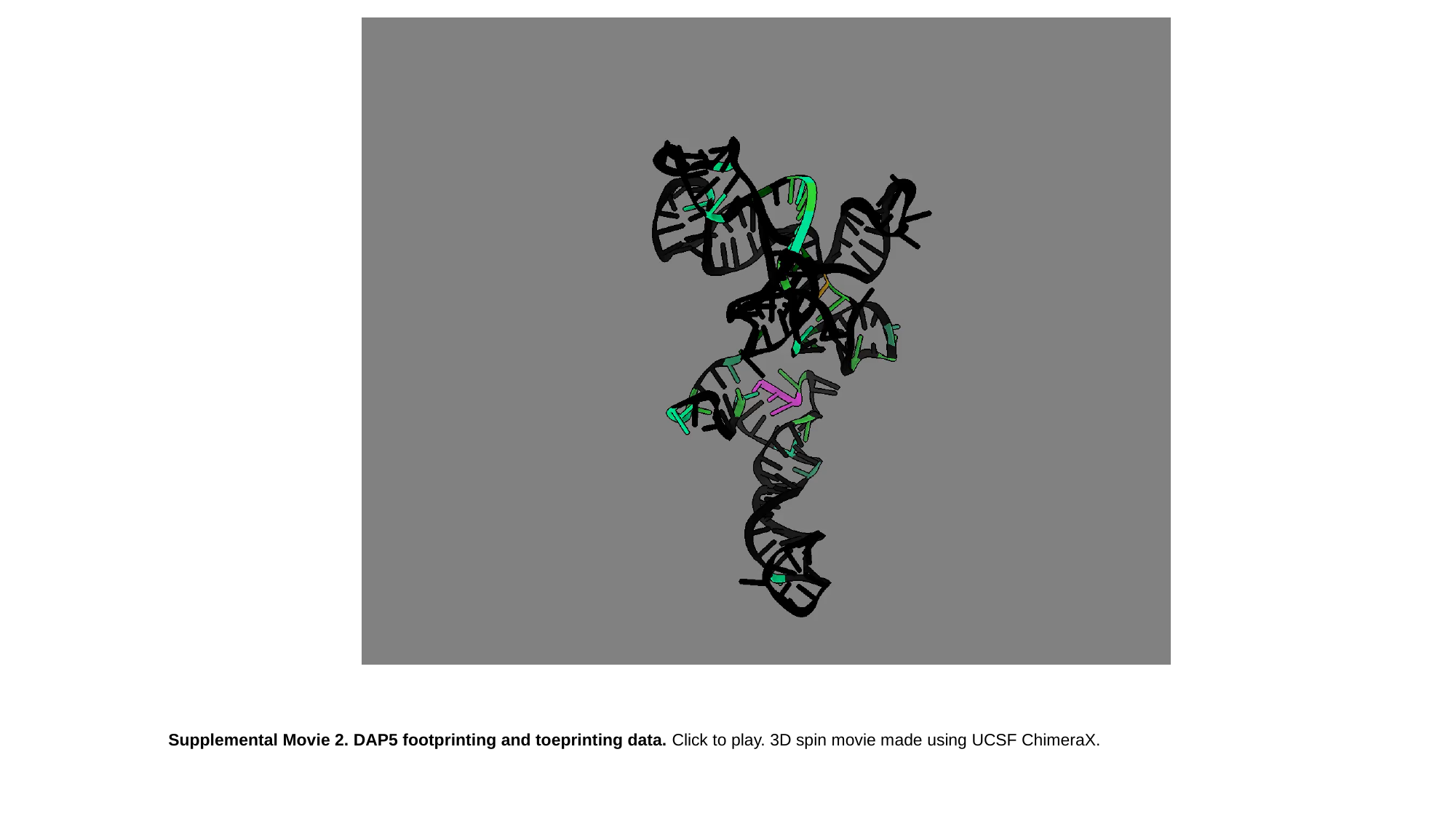

Supplemental Movie 2. DAP5 footprinting and toeprinting data. Click to play. 3D spin movie made using UCSF ChimeraX.

### Slide 7
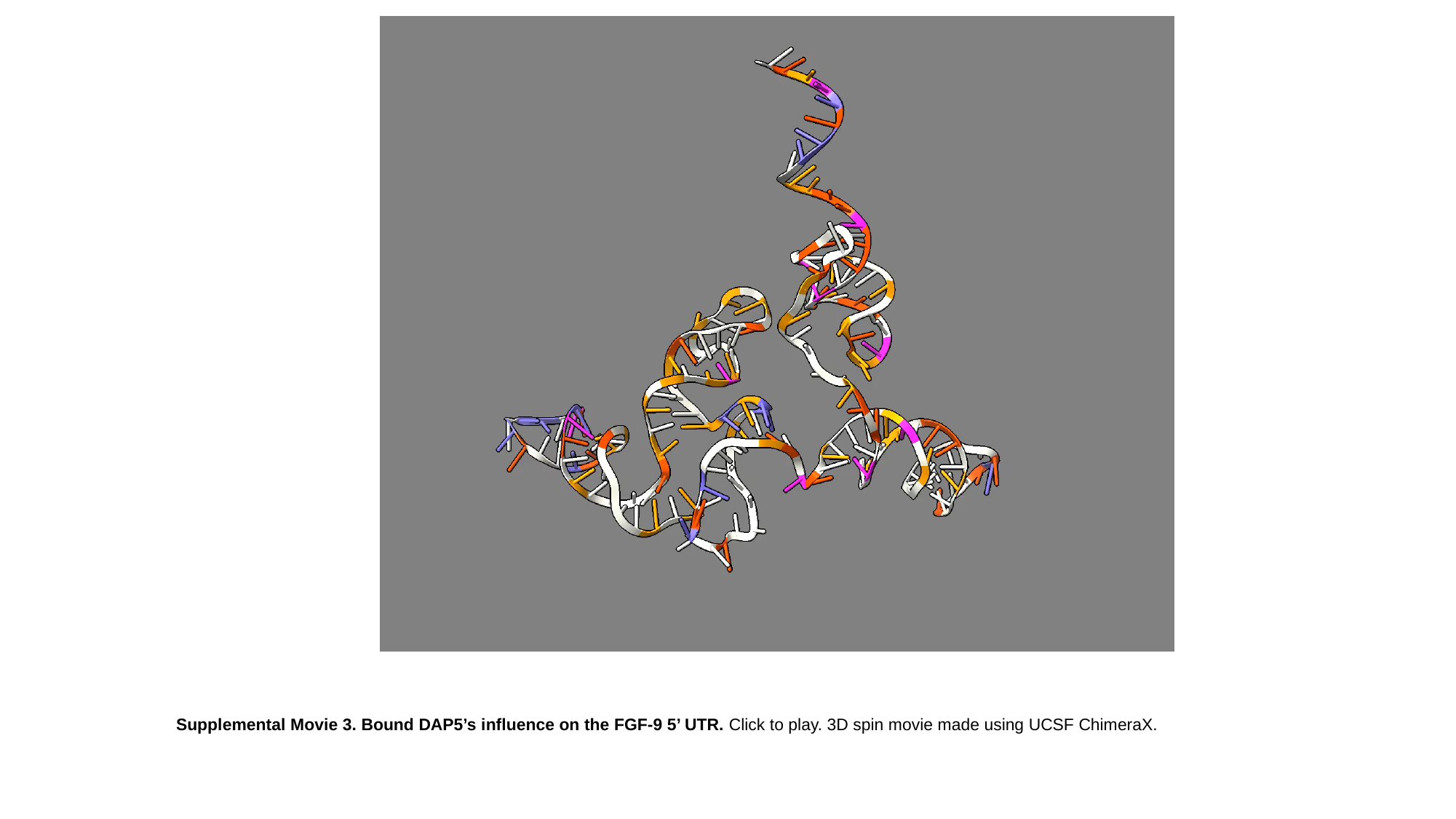

Supplemental Movie 3. Bound DAP5’s influence on the FGF-9 5’ UTR. Click to play. 3D spin movie made using UCSF ChimeraX.

### Slide 8
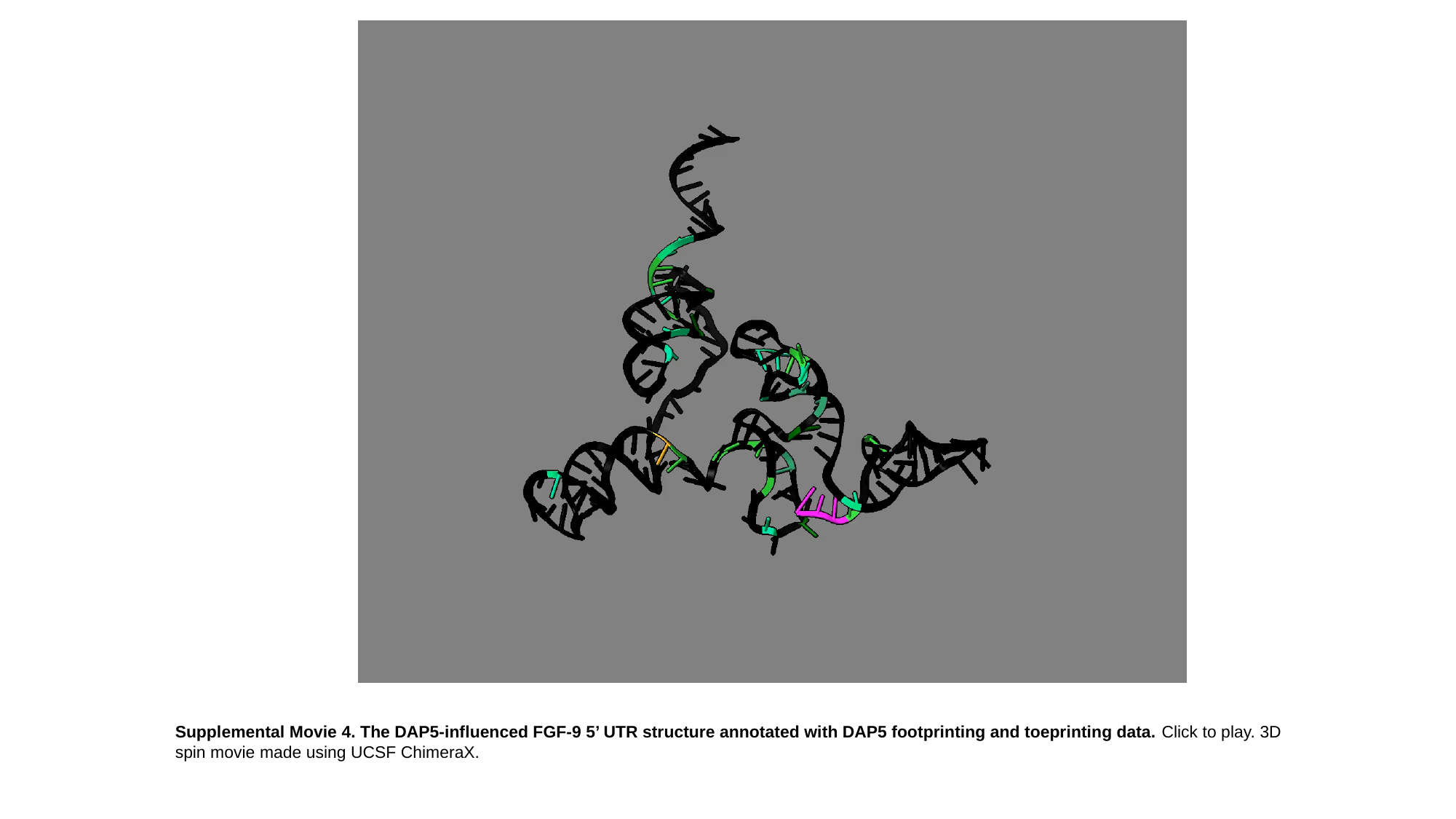

Supplemental Movie 4. The DAP5-influenced FGF-9 5’ UTR structure annotated with DAP5 footprinting and toeprinting data. Click to play. 3D spin movie made using UCSF ChimeraX.
